# Group 3 Innate Lymphoid Cells are regulated by WASP in a microbiota-dependent manner

**DOI:** 10.1101/2022.07.19.500438

**Authors:** Amlan Biswas, Naresh S Redhu, Anubhab Nandy, Yu Hui Kang, Michael Field, Ryan Kelly, Liza Konnikova, Jeremy A. Goettel, Amy M. Tsou, Bruce Horwitz, Scott B. Snapper

## Abstract

Wiskott-Aldrich syndrome protein (WASP) is a cytoskeletal regulator that is largely restricted to hematopoietic cells. While WASP expression in both lymphocytes and macrophages play a critical role in maintaining intestinal homeostasis, the function of WASP in innate lymphoid cells is unknown. Here we analyzed the role of WASP in the differentiation and function of group 3 innate lymphoid cells (ILC3s). WASP-deficient mice (*Was*^-/-^) have a marked reduction in ILC3s. Moreover, antimicrobial peptide expression in response to ILC3-derived IL-22 was also reduced in the absence of WASP. In *Was*^*-/-*^ mice, we observed a reduction in CCR6^+^ ILC3s, cells known to restrict immune responses to commensal bacteria. WASP-deficient mice were more susceptible to *Citrobacter rodentium*, an enteric infection controlled by ILC3s. Interestingly, there was no reduction in ILC3s in Was^-/-^ germ-free mice when compared to WT germ-free mice. ILC3s lacking *WASP* expression also demonstrated microbially-dependent alterations in gene expression associated with cell migration. Finally, ILC3-like (Rorgt^+^CD3^-^) cells were reduced in the GI tract of WASP-deficient patients. In conclusion, ILC3-specific expression of WASP is critical for the generation and function of ILC3s in the presence of commensal microflora. Aberrant ILC3 function in the setting of WASP-deficiency may contribute to underlying disease pathogenesis.

## Introduction

Wiskott–Aldrich syndrome protein (WASP) is a hematopoietic-specific member of a family of actin regulators that is defective in patients with mutations in the *WAS* gene. These patients suffer from recurrent infections, thrombocytopenia, eczema, and autoimmunity^1^. A subset of Wiskott-Aldrich syndrome (WAS) patients, similar to *Was*^*-/-*^ mice raised under specific pathogen free (SPF) conditions, also develop intestinal inflammation^3,4^. Over the last two decades, we and others have delineated functional roles for WASP in cell migration as well as immunoregulatory roles in lymphocytes and myeloid cells^2^. We have demonstrated that regulatory T cells and anti-inflammatory macrophages require WASP for maintaining immune tolerance^5,6^. WASP deficiency in macrophages leads to aberrant generation of anti-inflammatory macrophages and development of colitis^6^. Mechanistically, a WASP-DOCK8 complex regulates tolerogenic IL10-dependent signaling in macrophages. In addition, WASP has been reported to have regulatory affects in other myeloid cells including DCs, neutrophils, and mast cells^2^.

Numerous recent studies have identified innate lymphoid cells (ILCs) as critical mediators of intestinal homeostasis and immune tolerance. ILCs promote intestinal immune homeostasis through a variety of mechanisms including promoting barrier function, dampening commensal-specific T cell responses, and inducing regulatory T cells^7–11^. RORγt-expressing ILC3s are the most abundant ILC subset in the ileum and colon^12,13^. ILC3s mediate homeostatic control of intestinal immunity by expressing a range of cytokines including IL22 and IL17, cytokines known to induce expression of antimicrobial peptides and to maintain barrier integrity^14–16^. ILC3s are critical for the clearance of the intestinal pathogen *Citrobacter rodentium* as well as containment of commensal *Alcaligenes spp*^17^. Moreover, ILC3s regulate tolerogenic functions of DCs and macrophages^18^.

Here we investigated the role of WASP in the regulation of ILCs in the intestine. We found that WASP expression is critical for the maintenance and function of ILC3s in a microbiota-dependent manner. Both ubiquitous and ILC3-specific deletion of WASP resulted in enhanced susceptibility to *C. rodentium-*induced colitis. Similarly, WAS patients also exhibit a reduction in the number of ILC3s in the intestine. Collectively, these data demonstrate that WASP plays a critical role in ILC3-mediated regulation of immune tolerance.

## Results

### WASP deficiency results in reduced RORγt^+^ ILC3s in the gastrointestinal tract

ILCs form a major part of the immune cell repertoire at barrier mucosal surfaces and provide indispensable protection from numerous infectious microbes. We have previously shown that WASP deficiency in innate immune cells is a primary driver of intestinal inflammation^19^, and more recently demonstrated that WASP is important for anti-inflammatory macrophage function and mucosal immune tolerance^6^. However, whether ILCs require WASP for optimal development and intestinal mucosal-protective immunity remains poorly defined^20^. Therefore, we first examined the ILC compartment of WT and *Was*^*-/-*^ mice by staining lamina propria (LP) cells from both the small intestine and colon^21^. Frequencies of both group 1 (NKp46^+^) and group 2 (KLRG1^+^) ILCs were comparable between WT and *Was*^*-/-*^ mice (Supplementary Figure S1).

Interestingly, both the percentage and absolute numbers of ILC3s (RORγt^+^) were profoundly reduced in both the SI and colonic LP of *Was*^*-/-*^ mice (Figure 1A). CCR6^+^ ILC3s are abundant in the mLN of healthy mice and are known to selectively limit the expansion of commensal bacteria-specific effector CD4^+^ T cell responses^7^. In addition, CCR6^+^ ILC3s were significantly reduced in the mLN of *Was*^*-/-*^ mice (Figure 1B). Consistent with this defect, *Was*^*-/-*^ mice at 16 weeks of age exhibited robust levels of commensal bacteria-specific IgG in their serum compared to that of WT mice, which was statistically significant even at 6 weeks of age (Figure 1C).

**Figure 1:**
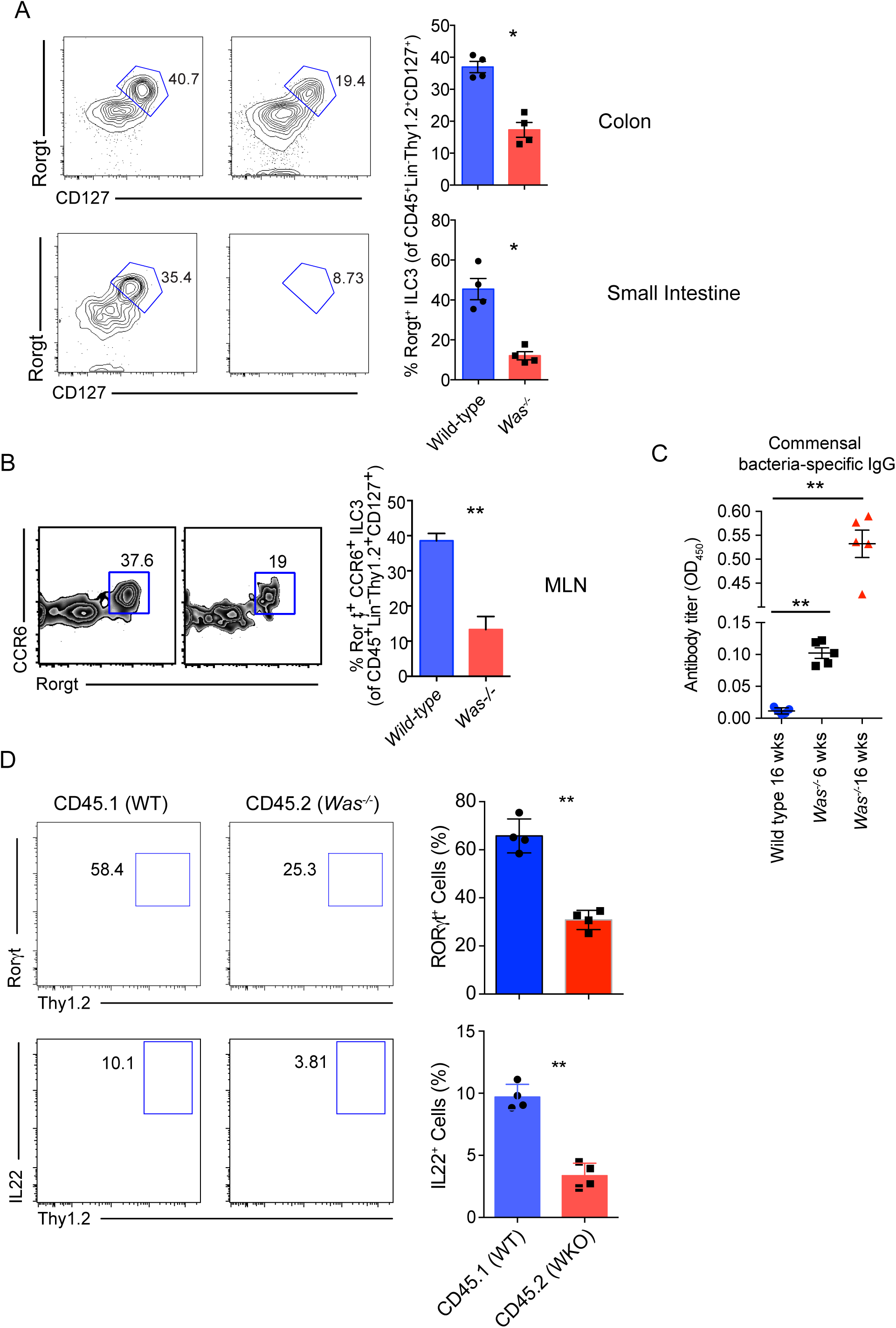
Reduced ILC3 abundance in the GI tract of *Was*^*-/-*^ mice. (A) Flow cytometric plots of ILC3s in colonic and small intestinal LP of indicated mice, followed by quantitation of frequencies on the right. ILC3s were gated as live CD45^+^Lin^-^Thy1.2^+^CD127^+^RORγt^+^ cells. WT (n=4) and *Was*^*-/-*^ (n=4) mice *p<0.05, unpaired *t* test. Representative of 2 independent experiments (B) Flow cytometric plot and quantitation of CCR6^+^ ILC3 in the mesenteric lymph nodes (mLN) of WT (n=3) and *Was*^*-/-*^ (n=3) mice. Representative of 2 independent experiments (C) Commensal bacteria-specific IgG titers in the serum of WT and *Was*^*-/-*^ mice determined by ELISA at indicated ages. Wild type 16 weeks (n=5), *Was*^-/-^ 6 weeks (n=5), *Was*^-/-^ 16 weeks (n=5). **p<0.01, unpaired *t* test. Representative of 2 independent experiments. (D) CD45.1^+^ (WT) and CD45.2^+^ (*Was*^*-/-*^) bone marrow cells were transferred at a ratio of 1:1 into lethally irradiated CD45.2^+^ *Was*^*-/-*^ recipients. LP ILC3s were analyzed after 10 weeks. FACS plot shows the gating strategy for RORγt^+^ ILC3s on top, IL22^+^ ILC3s on the bottom from CD45.1 and CD45.2 fractions of a mixed chimera recipient. Graphs on the right show the quantification of indicated cells in the WT and *Was*^*-/-*^ compartment of recipient mice. WT (n=4) and *Was*^*-/-*^(n=4), **p < 0.01, unpaired *t* test. Representative of 2 independent experiments.

To address whether the reduced frequency of ILC3s in *Was*^*-/-*^ mice was due to a cell-intrinsic defect caused by the absence of WASP, we performed radiation mixed bone marrow (BM) chimeras. BM from *Was*^*-/-*^ (CD45.2^+^) mice and congenic WT (CD45.1^+^) mice were injected intravenously at a 1:1 ratio into irradiated *Was*^*-/-*^ (CD45.2^+^) mice and analyzed 8 weeks after reconstitution. As shown in Figure 1D, *Was*^*-/-*^ BM progenitors gave rise to less RORγ ILC3s when compared to that generated from WT BM ((Figure 1D, top panel). To test whether these ILCs were functionally competent, we evaluated the production of IL-22 following IL-23 stimulation of LP cells from the mixed chimeras. Interestingly, IL-22^+^ ILC3s were profoundly reduced in the *Was*^*-/-*^ (CD45.2^+^) compartment (Figure 1D, bottom panel). Taken together, our data show that WASP is required for the development and/or maintenance of ILC3s in a cell-intrinsic manner.

ILC3s share multiple phenotypic and functional characteristics with T-helper 22 (Th22) cells as both cell types produce large amounts of IL22^22^. Therefore, we sought to confirm the requirement of WASP in ILC3 development in *Was*^*-/-*^ *Rag1*^*-/-*^ DKO mice, which lack both mature B and T cells. Consistent with *Was*^*-/-*^ mice, *Rag1*^*-/-*^*Was*^*-/-*^ mice showed significantly reduced colonic ILC3s compared to *Rag1*^*-/-*^ mice (Figure 2A). IL-22 production from ILC3s was also attenuated in *Rag1*^*-/-*^*Was*^*-/-*^ mice compared to *Rag1*^*-/-*^ mice (Figure 2B). These data confirm that ILC3s require WASP for their peripheral maintenance and effector cytokine production independent of adaptive immune cells.

**Figure 2:**
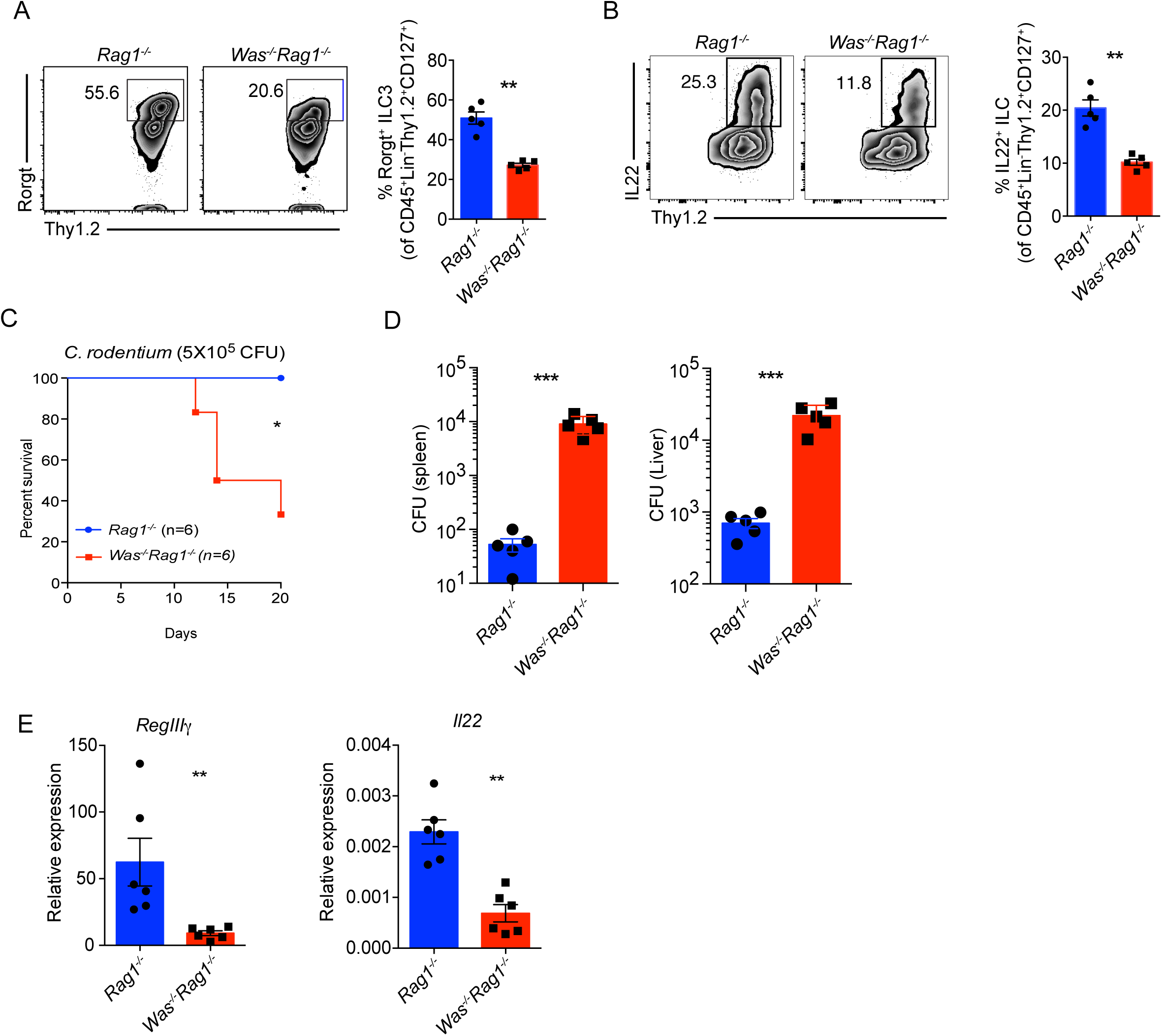
WASP expression in ILC3s is required for protective immunity against *C. rodentium*. Representative FACS plots and comparison of the frequencies of ILC3s (A), and IL-22^+^ ILC3s (B), gated as shown, in the colonic LP of *Rag1*^*-/-*^ and *Was*^*-/-*^*Rag1*^*-/-*^ mice. **p<0.01, unpaired *t* test. *Rag1*^*-/-*^ (n=5) and *Was*^*-/-*^ *Rag1*^*-/-*^ (n=5), Representative of 3 independent experiment. Percent survival (C) of *Rag1*^*-/-*^ and *Was*^*-/-*^*Rag1*^*-/-*^ mice following *C. rodentium* infection. *p<0.05, ****p<0.0001, unpaired *t* test. (D) CFUs of *C. rodentium* 3 weeks post-infection from the spleens and livers of indicated mice. ***p<0.001, unpaired *t* test. (E) Relative expression by qRT-PCR of *Reg3g* and *Il22* transcripts in the colonic tissue of *Rag1*^*-/-*^ (n=6) and *Was*^*-/-*^*Rag1*^*-/-*^ (n=6) mice 3 weeks post *C. rodentium* infection. Representative of 3 independent experiment.

### WASP expression in RORγt^+^ ILCs regulates protective immunity against *C. rodentium*

To assess whether WASP-deficient ILC3s regulate the mucosal immune response to an intestinal bacterial pathogen, we infected age-matched *Rag1*^*-/-*^ and *Rag1*^*-/-*^*Was*^*-/-*^ mice with a sublethal dose of *Citrobacter rodentium* by oral gavage. Compared to *Rag1*^*-/-*^ mice, which were fully protected, *Rag1*^*-/-*^*Was*^*-/-*^ mice succumbed to infection starting 12 days post-infection (Figure 2C). Furthermore, *C. rodentium* CFUs were markedly increased in the spleens and livers of *Was*^*-/-*^*Rag1*^*-/-*^ mice when compared to *Rag1*^*-/-*^ mice (Figure 2D), indicating that *Was*^*-/-*^*Rag1*^*-/-*^ mice were unable to contain bacterial dissemination. Since IL22-driven induction of the antimicrobial peptide RegIIIγ in colonic epithelial cells is required for protection from *C. rodentium*^23,24^, we evaluated the expression of these genes in the colonic tissue of *C. rodentium*-infected animals. As shown in Figure 2E, both *Reg3g* and *IL22* transcripts were substantially reduced in the colonic tissue of *Was*^*-/-*^*Rag1*^*-/-*^ mice compared to *Was*^*-/-*^ mice following *C. rodentium* infection.

To rule out the possibility that the reduction in ILC3s in *Was*^*-/-*^*Rag1*^*-/-*^ mice resulted from WASP deficiency in other innate immune cells, we assessed ILC3s in mice conditionally targeted for WASP in RORγt+ cells. As shown in Figure 3A, both the colon and SI LP showed significantly reduced levels of ILCs in *Was*^*fl/fl*^*Rorc*^*Cre*^*Rag1*^*-/-*^ mice compared to *Was*^*fl/fl*^*Rag1*^*-/-*^ mice. IL-22 production from ILC3s was also reduced in *Was*^*fl/fl*^*Rorc*^*Cre*^*Rag1*^*-/-*^ mice compared to *Was*^*fl/fl*^*Rag1*^*-/-*^ controls in both the colon and SI LP (Figure 3B). Finally, we assessed whether conditionally targeted WASP-deficient mice also had increased susceptibility to *C. rodentium* infection. When compared to *Was*^*fl/fl*^*Rag1*^*-/-*^ mice, *Was*^*fl/fl*^*Rorc*^*Cre*^*Rag1*^*-/-*^ mice had enhanced mortality and systemic bacterial dissemination, upon *C. rodentium* infection (Figure 3C, D). Taken together, these data indicate that WASP is necessary for RORγt^+^ ILC3-dependent protection against an enteric pathogen.

**Figure 3:**
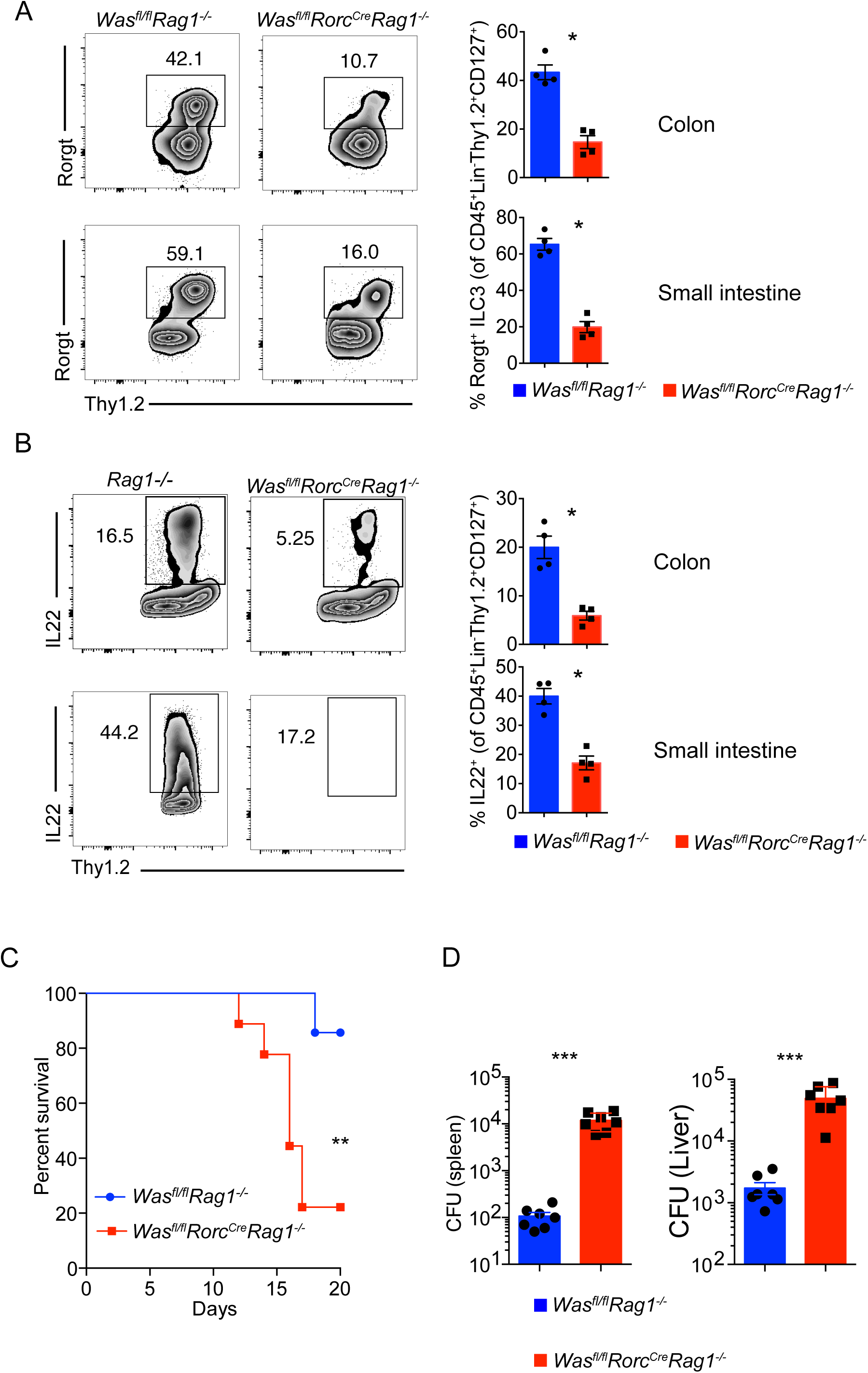
WASP expression in ILC3s is necessary for protection from *C. rodentium* infection. Representative FACS plots and comparison of the frequencies (right) of ILC3s (A), and IL-22^+^ ILC3s (B), gated as shown, in the colonic and SI LP of *Was*^*fl/*^ *Rag1*^*-/-*^ (n=4) and *Was*^*fl/fl*^*Rorc*^*Cre*^*Rag1*^*-/-*^ (n=4) mice. *p<0.05, unpaired *t* test. Representative of 3 independent experiments. Percent survival (C), and CFUs of *C. rodentium* in the spleens and livers (D) of *Rag1*^*-/-*^ (n=7) and *Was*^*fl/fl*^*Rorc*^*Cre*^*Rag1*^*-/-*^ (n=7) mice 3 weeks post-infection with *C. rodentium*. **p<0.01, ****p<0.0001, unpaired *t* test. Representative of 2 independent experiment.

### ILC3 homeostasis in WASP-deficient mice is microbiota-dependent

ILC3s have been shown to regulate selective anatomical containment of lymphoid-resident commensal microbiota^17^, and reciprocally, multiple microbiota-derived signals have been shown to control the differentiation and function of ILC3s^25^. To assess whether the role of WASP in regulating ILC3 differentiation/development is dependent on intestinal microbiota, we treated *Rag1*^*-/-*^ and *Was*^*-/-*^*Rag1*^*-/-*^ mice with broad-spectrum antibiotics and analyzed the ILC3 compartment 4 weeks post antibiotic treatment. As shown in Figure 4A, the frequencies of RORγt^+^ and IL-22-producing ILC3s were comparable in Abx-treated *Was*^*-/-*^*Rag1*^*-/-*^ and *Rag1*^*-/-*^ mice (Figure 4A).

**Figure 4:**
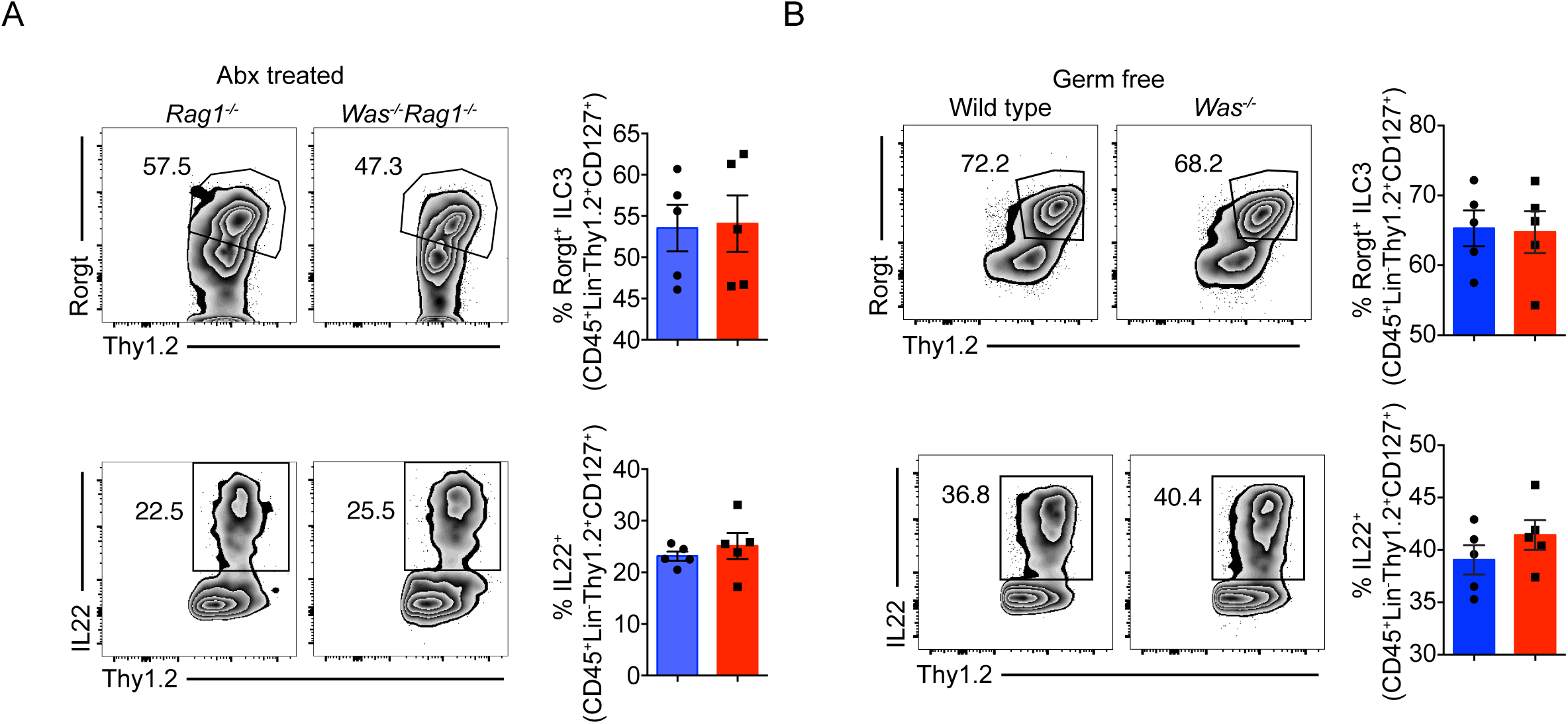
Microbiota regulates ILC3 phenotype in the colonic LP of WASP-deficient mice. (A) Representative FACS plots and comparison of the frequencies (right) of RORγt^+^ ILC3s (top), and IL-22^+^ ILC3s (bottom), gated as shown, in the colonic LP of *Rag1*^*-/-*^ (n=5) and *Was*^*-/-*^*Rag1*^*-/-*^ (n=5) mice following broad spectrum antibiotic (Abx) treatment for 3 weeks. Representative of 2 independent experiments. (B) Representative FACS plots of RORγt^+^ ILC3s (top), IL-22^+^ ILC3s (bottom), and respective frequencies (right) in the colonic LP of germ-free (GF) WT (n=5) and *Was*^*-/-*^ (129SvEv) mice (n=5). *p<0.05, unpaired *t* test. Representative of 2 independent experiments.

We further assessed the role of commensal microbiota in regulating ILC3 development in germ-free *Was*^*-/-*^ mice. Consistent with antibiotic-treated mice, germ-free *Was*^*-/-*^ mice exhibited similar levels of RORγt^+^ and IL-22-producing ILC3s as germ-free WT mice (Figure 4B). Collectively, these data indicate that the microbiota regulate the ILC3 compartment in *Was*^*-/-*^ mice.

### ILC3 of *Was*^*-/-*^ mice exhibit altered gene expression profile, associated with cell migration

WASP is necessary for cytoskeleton-dependent functions including podosome formation and cell migration^26–31^. Since we observed a marked reduction in ILC3 in the colons of *Was*^*-/-*^ mice in the SPF but not in the germ-free setting, we hypothesized that a microbiota-dependent cellular function is compromised in WASP deficiency. To examine this, we performed bulk RNA sequencing (RNAseq) on FACS-sorted colonic ILC3s from SPF *Was*^*-/-*^ *Rag1*^*-/-*^ and *Rag1*^*-/-*^ mice (C57BL/6 background), and germ free *Was*^*-/-*^ and WT mice (129SvEv background). We identified a total of 513 differentially expressed genes (padj <0.05) between ILC3s from *Was*^*-/-*^*Rag1*^*-/-*^ and *Rag1*^*-/-*^ mice in SPF conditions (Supplementary table S1). Some of these genes (Figure 5A) such as *Fbln1, Plekhh2, Pcdh18*, and *Sparc* are known to regulate cell adhesion, motility, and migration in various cell types^31–34^. Consistently, gene ontology (GO) analysis of the 513 differentially expressed genes in ILC3s from *Was*^*-/-*^ *Rag1*^*-/-*^ and *Rag1*^*-/-*^ mice revealed categories related to cell motility, locomotion, chemotaxis, and cell migration (Figure 5B). Interestingly, however, the RNAseq data obtained from ILC3s isolated from germ-free *Was*^*-/-*^ and WT mice revealed only 25 differentially expressed genes (padj<0.05) (Supplementary table S2, Figure 5C), suggesting that the microbiota is a major driver of the transcriptional signature observed in SPF mice. Taken together, these data suggest that ILC3s isolated from the colons of WASP-deficient mice exhibit transcriptional signatures associated with defect(s) in cell migration/recruitment in a microbiota-dependent manner.

**Figure 5:**
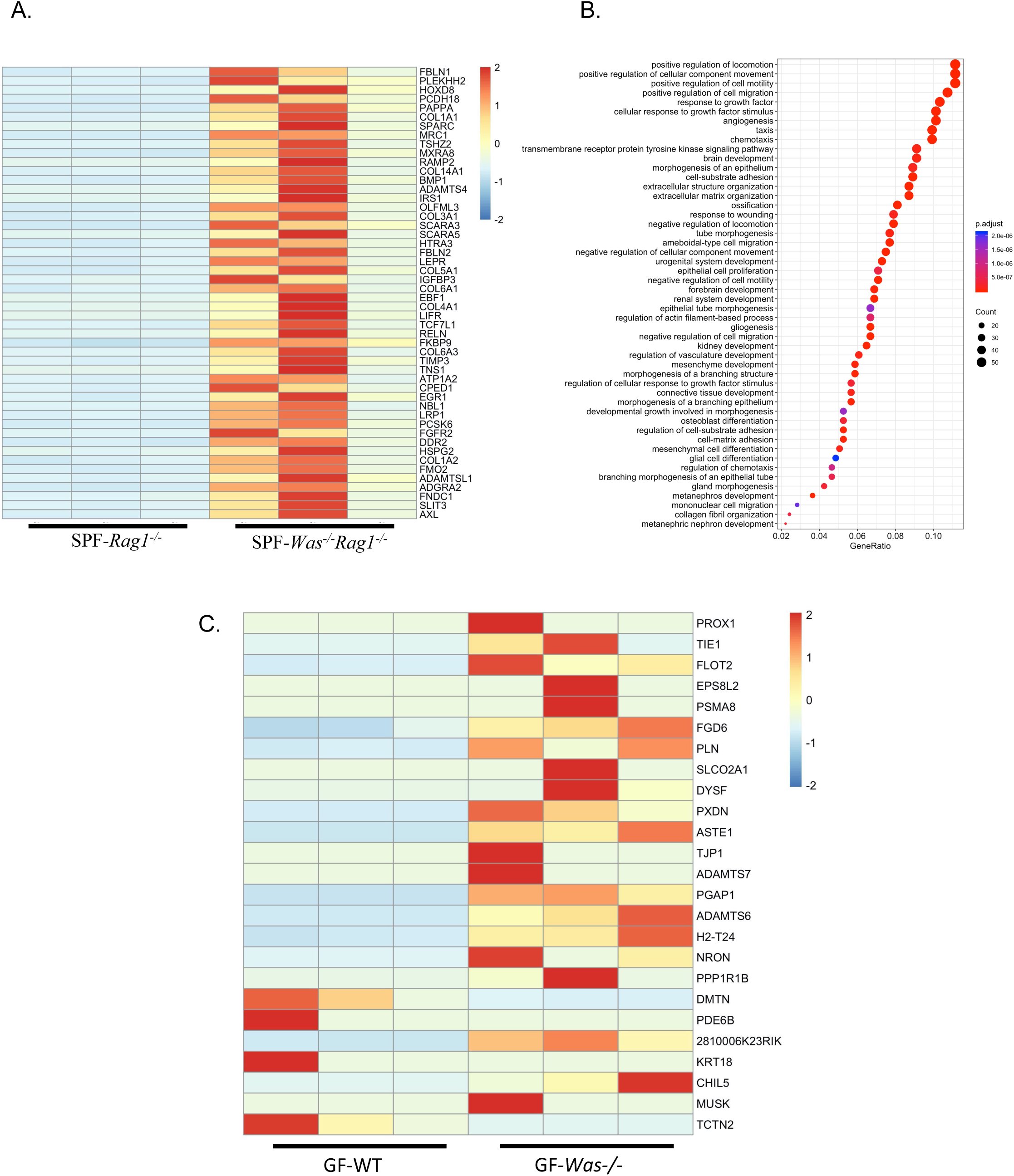
ILC3 of WASP-deficient mice exhibit microbiota-dependent defective cell migration transcriptional profile. Colonic LP ILC3s (gated on Lin^-^Thy1.2^hi^CD45^lo^ cells) were sorted (10-100×10^3^ cells), RNA was prepared, and RNAseq was performed. (A) Heatmap showing relative expression intensity of top 50 differentially expressed genes in the LP ILC3s between specific pathogen-free (SPF) *Rag1*^*-/-*^ and *Was*^*-/-*^*Rag1*^*-/-*^ mice. (B) Gene ontology (GO) analysis of total 513 differentially expressed genes (padj<0.05) from samples processed for RNAseq in (A). (C) Top 25 differentially expressed genes (padj<0.05) in the LP ILC3 between germ-free (GF) WT and *Was*^*-/-*^ mice.

### Intestinal RORγt^+^ ILC-like cells are reduced in patients with WASP deficiency

Finally, we sought to investigate whether patients with WAS also exhibit defects in ILC3s like *Was*^*-/-*^ mice. To this end, we performed RNA in situ hybridization (RNAscope) on formalin fixed paraffin embedded colonic biopsy tissues. As shown in Figure 6A, we identified cells positive for transcripts encoding CD3 and RORγt, marked as CD3^+^, CD3^+^RORγt^+^, and CD3^-^RORγt^+^ cells. The intestinal tissue from five out of six WAS patients showed significantly reduced abundance for CD3^-^RORγt^+^ cells compared to matched controls (Figure 6B). Collectively, our data suggest that WASP deficiency is associated with reduction in ILC3s in the GI tract of both mice and humans.

**Figure 6:**
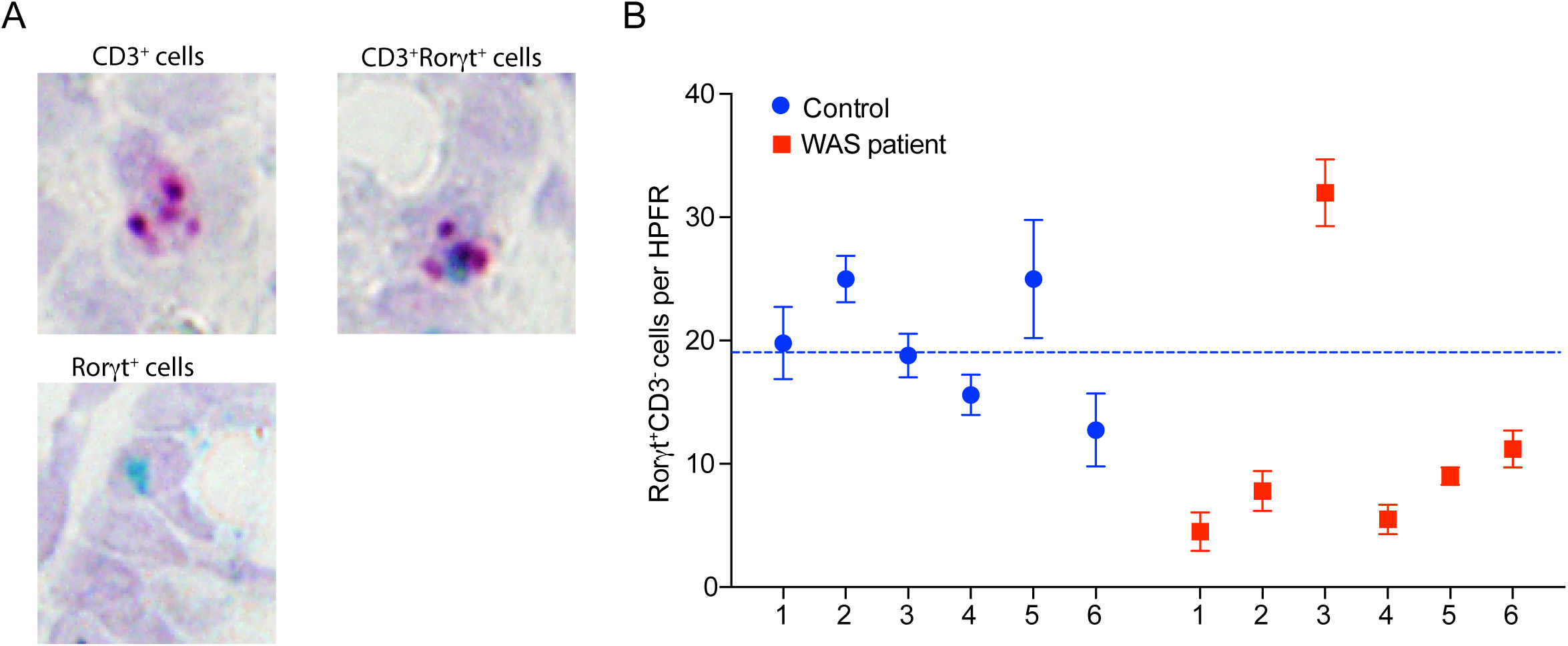
ILC3-like RORγt^+^CD3^-^ cells are reduced in the GI tract of WAS patients. (A) FFPE intestinal biopsy tissue sections showing RNA in situ hybridization (ISH) performed by RNAScope® using CD3 and RORγt probes. (B) Quantitation of RORγt^+^CD3^-^ cells per high power field in the tissue sections stained in (A) from control and WAS patients.

## Discussion

ILCs are critical regulators of mucosal homeostasis that play a crucial role in the maintenance of epithelial barrier integrity, control of pathogen dissemination and amelioration of intestinal injury^35,36^. Among the different subsets of ILCs, ILC3s are present in the greatest abundance in the intestine, and their numbers depend on microbial cues^21,25^. ILC3s maintain intestinal mucosal immune homeostasis by regulating a plethora of immune functions including the production of antimicrobial peptides from intestinal epithelial cells, suppression of commensal specific T-cell responses, generation/maintenance of Tregs, and the induction of antibody production from both mucosal and splenic B cells^7,8,11,37^. Alterations in intestinal ILC3 function, resulting from genetic or environmental perturbations, can lead to aberrant immune activation and associated intestinal inflammation^38^. WASP expression in hematopoietic cells is critical for intestinal homeostasis, including the regulation of intestinal anti-inflammatory macrophages, suppression of pathogenic Th2 responses and induction of IL10 production by Tregs^2,4,5,6,18^. A role for WASP expression in ILC-dependent regulation of mucosal homeostasis had not been explored previously.

Our findings demonstrate a critical role for WASP in regulating ILC3s. WASP deficiency in mice resulted in reduced ILC3 frequency in the intestine associated with enhanced susceptibility to the enteric pathogen *C. rodentium*. Selective deletion of WASP in ILC3s was also associated with reduced ILC3s and enhanced *C. rodentium*-mediated colitis, demonstrating a cell-intrinsic role for WASP in ILC3-dependent functions. Moreover, RNAscope analysis in colonic tissue biopsies of WAS patients revealed a reduction in ILC3 numbers. Overall, our data indicate that WASP is a critical regulator of ILC3s in the GI tract of both mice and humans.

Microbial factors are essential for the distribution and maintenance of ILC3s^39^. The maternal microbiota contributes to the maintenance of ILC3s, as pups obtained from transiently colonized dams have higher intestinal NKp46^+^RORγt^+^ ILC3s compared to germ-free dams^40^. Recent advances suggest that microbial factors regulate cell adhesion and interactions with the extracellular matrix^21^. Interestingly, while *Was*^*-/-*^ mice housed in SPF conditions have reduced numbers of ILC3s, germ-free or antibiotic-treated *Was*^*-/-*^ mice display no significant alteration in ILC3s compared to germ-free or antibiotic-treated WT mice. Differentially expressed genes in ILC3s from *Was*^*-/-*^*Rag1*^*-/-*^ compared to *Rag1*^*-/-*^ mice, which are primarily associated with cell migration and adhesion, were not significantly different between ILC3s from germ-free *Was*^*-/-*^ versus WT mice. Taken together, these data suggest that ILC3 homeostasis, ILC3 migration and/or cell retention, is WASP-dependent only in the presence of the gut microbiota.

In summary, we have discovered a novel function of WASP in the microbially-dependent regulation of ILC3 maintenance in the GI tract. Our data indicate that reduction in RORγt^+^ ILC3s and defective IL-22 production by these cells result in increased mortality of WASP-deficient mice after *C. rodentium* infection. Reduced numbers of CD3^-^RORγt^+^ cells in the mucosa of WAS patients strengthen the significance of these murine findings. Given that WAS patients exhibit recurrent infections and a subset of these patients develop IBD, defective RORγt^+^ ILC3s may contribute to these mucosal findings. Collectively, our work clearly highlights the critical role for WASP in regulating mucosal homeostasis through maintenance of the ILC3 population in the gut. We believe that our observation of microbiota-dependent WASP-mediated ILC3 regulation in the gut emphasizes the need for understanding the complex relationship between commensal microbiota and host immune cells at the mucosal interface.

## Methods

### Study approval

All patients provided informed consent prior to inclusion in the study. Clinical patient samples were collected under a Boston Children’s Hospital IRB-approved research protocol (P00000529). All animal experiments were performed in accordance with Institutional Animal Care and Use Committee-approved protocols number 14-04-2677R (IACUC, Boston Children’s Hospital), and adhered to the National Research Council’s ‘Guide to the care and Use of Laboratory Animals.

### Mice

C57BL/6 background *Was*^*fl/fl*^ mice have been described previously and were crossed with *Rorc*^*Cre*^ (Cat. No. 022791) mice obtained from The Jackson Laboratories to generate *Was*^*fl/fl*^*Rorc*^*Cre*^. *Was*^*fl/fl*^*Rorc*^*Cre*^ mice were subsequently crossed with *Rag1*^*-/-*^ mice to generate *Was*^*fl/fl*^*Rorc*^*Cre*^*Rag1*^*-/-*^ to study the intrinsic role of WASP in ILC3. *Was*^*-/-*^, *Was*^*-/-*^*Rag1*^*-/-*^ were generated as previously described^20^. All the mice were housed in micro-isolator cages in a specific pathogen-free animal facility at Boston Children’s Hospital. Sex-and age-matched animals between 5 and 14 weeks of age were used for experiments. Germ free WT and *Was*^*-/-*^ mice in 129SvEv background were housed in germ-free (GF) facility at Boston Children’s Hospital. GF mice were maintained in sterile isolators and fed autoclaved food and water. Mice were screened weekly for contamination by bacterial plating and PCR. We did not use randomization to assign animals to experimental groups. Investigators were not blinded to group allocation during experiments. No animals were excluded from the analysis. Experiments were conducted after approval from the Animal Resources at Children’s Hospital and according to regulations of the Institutional Animal Care and Use Committees (IACUC).

### Reagents and antibodies

The antibiotics vancomycin, polymyxin B, ampicillin, and neomycin were obtained from Gold Biotechnology, Inc. (St. Louis, MO), and metronidazole was from Sigma-Aldrich Corp. (St. Louis, MO). The following reagents and anti-mouse antibodies were used for flow cytometry: Zombie Violet^™^ fixable viability kit (BioLegend, San Diego, CA), CD45 (clone 30-F11, BioLegend), CD11b (clone M1/70, BioLegend), CD11c (clone N418, BioLegend), NK1.1 (PK136, BioLegend), B220(RA3-6B2, BioLegend), CD3e (clone 145-2C11, BioLegend), CD4 (clone GK1.5, BioLegend), IL17A (clone TC11-18H10.1, BioLegend), IL22 (Poly5164, BioLegend), CD127 (clone A7R34, BioLegend), CD90.2 (Thy1.2, clone 30-H12, BioLegend), CD335 (NKp46, clone 29A1.4, BioLegend), KLRG1 (clone 2F1/KLRG1, BioLegend), and RORγt (clone B2D, eBioscience/ Thermo Fisher Scientific, Waltham, MA). The lineage marker cocktail (Biolegend) contains clones 145-2C11, M1/70, RA3-6B2; TER-119 and RB6-8C5.

### Isolation of LP cells

LP immune cells were prepared as we reported recently^6^. Briefly, SI and colons were stripped of epithelial cells by agitating in Hank’s balanced salt solution (HBSS, without Ca/Mg salts) containing 2% fetal bovine serum (FBS), 10mM EDTA, 1mM dithiothreitol and 5mM HEPES at 37°C for 30 minutes. After removal of the epithelial layer, tissues were washed in PBS and digested in buffer containing HBSS (containing Ca/Mg salts), 20% FBS and collagenase VIII (200 unit/ml) for 60 min. Undigested tissues were disrupted by repeated flushing through a 10ml syringe and the resulting single cell suspensions were filtered and stained for flow cytometry. The LP ILCs were gated as type 1 (Lin^-^ CD45^+^CD127^+^Thy1.2^+^ RORγt^-^NKp46^+^), type 2 (Lin^-^ CD45^+^CD127^+^Thy1.2^+^ RORγt^-^KLRG1^+^), and type 3 (Lin^-^CD45^+^CD127^+^Thy1.2^+^ RORγt^+^), as described by Gury-BenAri et al (2016)^21^.

### Flow cytometric analysis

LP cells were acquired using BD FACS Canto II flow cytometer or BD LSR Fortessa^™^ (BD Biosciences, San Jose, CA) and analyzed using FlowJo v9 and v10 (Treestar, Ashlan, OR). Anti-mouse CD16/32 was used as an Fc blocking reagent. For intracellular IL22 detection, LP cells were stimulated overnight in the presence of 15ng/ml recombinant mouse IL23 (Peprotech), followed by additional 4.5 hours with 500 ng/mL ionomycin and 20 ng/mL PMA in the presence of 1μg/mL of GolgiStop containing Monensin (BD Biosciences). Cell were then fixed, permeabilized (BD Biosciences), and stained for intracellular proteins.

### Generation of bone marrow chimera mice

CD45.2 *Was*^*-/-*^ recipient mice were irradiated with 1000 rad in 2 doses of 500 rad each 4 hours apart. Bone marrow cells from both femurs and tibiae of B6.SJL (CD45.1) and *Was*^*-/-*^ (CD45.2) donor mice were collected under sterile conditions and suspended in PBS before mixing at a 1:1 ratio and were then injected intravenously into each recipient mouse. The ratio of the bone marrow cells was confirmed by flow cytometry. Recipient mice were housed under specific pathogen-free conditions and were provided autoclaved water with sulfatrim (trimethoprim-sulfamethoxazole) and fed autoclaved food for three weeks. After three weeks they were provided normal food and water. Eight to ten weeks after the injection, ILC population in the colonic LP were examined.

### *Citrobacter rodentium* infection and CFU assessment

*C. rodentium* was cultured overnight at 37°C in Luria broth with gentle agitation. After spinning down at 4,500 r.p.m., they were suspended in PBS and the concentration was measured at A_600_. Mice were fasted for 8h before oral gavage of *C. rodentium* in 200 ul PBS. Colony-forming units from spleens and livers of infected mice were analyzed following mechanical dissociation in sterile PBS and after serial dilution, plated on LB agar plates and incubated overnight at 37°C.

### Quantitative RT-PCR

RNA from colonic tissue was isolated using TRIzol reagent (Life Technologies), and complementary DNA was reverse transcribed from 1 μg total RNA using iScript cDNA Synthesis Kit (Bio-Rad). mRNA expression was measured using iQ SYBR Green on a CFX96 Real-Time System (Bio-Rad). Expression of the transcripts was normalized against hypoxanthine-guanine phosphoribosyl transferase (HPRT) and presented as relative expression (2^-dCt^).

### Microbiota depletion of *Was*^*-/-*^ and wild-type mice

Intestinal microbiota in WT and *Was*^*-/-*^ mice were depleted by administrating cocktail of antibiotics as described previously. Briefly mice were treated with ampicillin (1 g/L), vancomycin (500 mg/L), neomycin sulfate (1 g/L), and metronidazole (1 g/L) in drinking water for three weeks. Efficiency of microbiota depletion was examined through the quantitative analysis of Eubacteria, Bacteroidetes and Firmicutes in the feces by qPCR using specific primers. After three weeks of antibiotic treatment, ILC population in the colonic LP of the mice were examined.

### RNA in situ hybridization (ISH) in tissue biopsies

Sections from formalin fixed paraffin embedded samples (control, n=5, and *WAS*, n=5, subjects) were obtained and freshly cut to 0.4-micron thickness as previously described by Wang et al^42^. Staining was completed per manufacturer’s instructions for RNAScope® 2.5 HD (chromogenic) duplex detection kit (Advanced Cell Diagnostics, ACD Bio, Newark, CA, USA). ROR*γ*t probe was used in C1 and CD3 probe was used in C2. All probes and reagents were obtained from ACD Bio. Images were taken at 40x with 6 images per patient and manual counting performed in ImageJ. The number of ROR*γ*t^+^ and CD3^-^ cells were counted per high power field and averaged per patient.

### ELISA to detect microbiota specific serum antibody

Commensal bacterial antigen was generated as described by Hepworth et al. Fecal contents from WT mice were homogenized and centrifuged at 1,000 rpm to remove large aggregates, and the resulting homogenate was washed with sterile PBS twice by centrifuging for 1 min at 8,000 rpm. On the last wash, bacteria were suspended in 2 mL ice-cold PBS and sonicated on ice. Samples were then centrifuged at 20,000g for 10 min, and supernatants were recovered for a crude commensal bacteria antigen preparation. For measurement of serum antibodies by ELISA, 5 µg/mL commensal bacteria antigen was coated on 96-well plates. Plates were subsequently washed 6 times with 300 μl of 0.05% Tween in PBS (washing buffer) and blocked with 10% fetal calf serum (FCS) in PBS for 2 hours at room temperature. Serum dilutions ranging from 1:30 to 1:5,000 were prepared in 2% FCS in PBS and transferred to wells in a volume of 100 ul. Plates were incubated at room temperature for 3 hours and then washed 6 times in washing buffer. Anti-commensal Ig antibodies were detected with horseradish peroxidase-conjugated antibodies against mouse Ig (Southern Biotech). Plates were developed in the dark with 100 ul of tetramethylenbenzidine per well (KPL). This reaction was stopped by the addition of 50 ul of 2M H_2_SO_4_ prior to spectrophotometric analysis at 450 nm (PerkinElmer).

### RNA sequencing and computational analysis

10,000–100,000 ILC3s, gated as Lin^-^CD90.2^hi^CD45^lo^ cells from colon LP were sorted on a FACS Melody into 350 ul of RLT Lysis buffer (Qiagen) and mRNA was isolated using RNeasy Plus mini kit (Qiagen). Oligo dT beads were used to enrich for mature polyA(+) transcripts. RNA-Seq was carried out at the Molecular Biology Core Facility (MBCF) of Dana-Farber Cancer Institute, Boston using the Nextseq Standard Program SE75, and reads were aligned to mm10 using TopHat with default settings, and DESeq was used to generate normalized counts for each gene. Only the genes that had normalized counts > 50 in at least one of the total six groups in SPF or GF subsets were further analyzed. Heatmaps were generated using ‘R’ studio from genes that were differentially expressed (top 50) between LP ILC3s of *Rag1*^*-/-*^ and *Was*^*-/-*^*Rag1*^*-/-*^ mice and top 25 between LP ILC3s of WT and Was-/-mice.

### Statistical analysis

All data were analyzed by unpaired *t* test with 95% confidence interval or ANOVA using GraphPad Prism version 6.0 (GraphPad software) and presented as mean±SEM. Normal distribution was assumed. A *p* value of less than 0.05 was considered statistically significant. (* P < 0.05; ** P < 0.01 and *** P < 0.001 and **** P < 0.0001; ns, not significant). All data shown is representative or cumulative of at least two, mostly three or more, independent experiments.

## Acknowledgements

This work was supported by the National Institute of Health Grants [P30 DK034854; 5P01 HL059561], the Helmsley Charitable Trust, the Wolpow Family Chair in IBD Treatment and Research, and the Boston Children’s Hospital Translational Investigator Service (to S. B. S), NIH KO1 award (K01DK109026) to A.B, Crohn’s, and Colitis Foundation of America RFA381023 to N.S.R, National Institute of Diabetes and Digestive and Kidney Diseases (NIDDK) DK106311 to J.A.G., Crohn’s and Colitis Foundation Research Fellowship Award (527125) to A.M.T.

## Author Contributions

AB- Conceptualization, Data curation, Formal analysis, Investigation, Writing—original draft, Writing—review and editing.

NSR- Data curation, Formal analysis, Investigation, writing—original draft, writing—review and editing.

AN- Investigation, writing—original draft, writing—review and editing.

MF- Investigation, Formal analysis JAG- Formal analysis, Investigation

AT- Formal analysis, Investigation

YHK- Formal analysis, Investigation, Data curation

RK- Sample Collection, Data curation

SBS- Conceptualization, Resources, Formal analysis, Funding acquisition, Investigation, Methodology, Writing—review and editing.

