## Supplemental Figure S1 for "Group 3 Innate Lymphoid Cells are regulated by WASP in a microbiota-dependent manner"

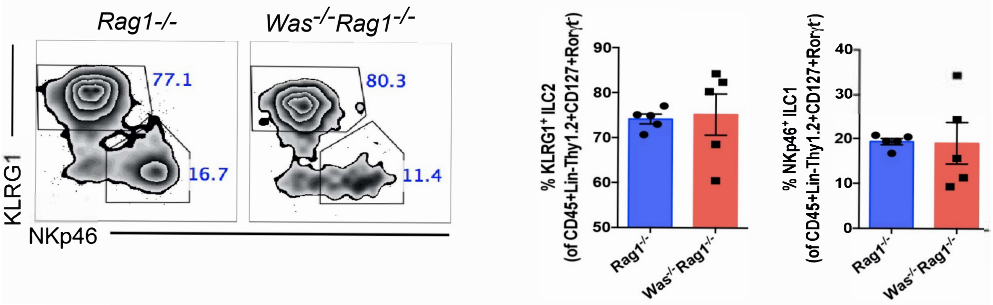


**Supplementary Figure S1: Normal ILC1 and ILC2 development in the colon of *Was*$^{-/-}$ mice.** Flow cytometric analysis of LP ILC1s and ILC2s (left) in mice at 12 weeks of age, followed by quantification (right) of ILC1s and ILC2s. ILC1s were gated as live CD45$^+$Lin$^-$Thy1.2$^+$CD127$^+$RORγt$^-$NKp46$^+$ whereas ILC2s were gated as live CD45$^+$Lin$^-$Thy1.2$^+$CD127$^+$RORγt$^-$KLRG1$^+$. Each dot in the right graphs represents an individual animal.
